# R-Ras coordinates reciprocal activation of ERK5 and ERK1/2 under single pathway inhibition in melanoma

**DOI:** 10.64898/2026.07.17.737152

**Authors:** Ignazia Tusa, Cosimo Mazzei, Dimitri Papini, Alessio Menconi, Ylenia Sfragano, Alessandro Tubita, Giulia Montemurro, Julia Penitenti, Azucena Esparis-Ogando, Atanasio Pandiella, Elisabetta Rovida

## Abstract

Malignant melanoma is frequently driven by constitutive activation of the RAS-RAF-MEK1/2-ERK1/2 pathway, yet adaptive signaling limits the long-term efficacy of MAPK-targeted therapies. Although activation of the MEK5-ERK5 pathway has emerged as a mechanism of resistance to RAF-MEK1/2-ERK1/2 inhibition, whether ERK5 inhibition reciprocally activates the canonical MAPK cascade and the molecular basis of this crosstalk remain unknown.

Here, we show that genetic and pharmacological inhibition of ERK5 induces further activation of the MEK1/2-ERK1/2 pathway in BRAFV600E melanoma cells. Based on our previous transcriptomic analyses, we investigated the role of the small GTPase R-Ras, identified among the genes upregulated following ERK5 silencing. Accordingly, R-Ras mRNA and protein levels increased upon both genetic and pharmacological ERK5 inhibition, whereas R-Ras silencing abolished ERK1/2 hyperactivation and potentiated the anti-proliferative and pro-apoptotic effects of ERK5 targeting. Conversely, inhibition of the RAF-MEK1/2-ERK1/2 pathway increased R-Ras expression and ERK5 activation, both of which were prevented by R-Ras depletion. Besides ERK1/2, overexpression of a constitutively active mutant of R-Ras promoted ERK5 activation, placing R-Ras upstream of both signaling cascades. Finally, the pan-Ras inhibitor RMC-6236 potentiated the antitumor activity of either ERK5- or RAF-MEK1/2-ERK1/2-targeted therapies in either two-dimensional cultures or melanoma spheroids.

Collectively, these findings identify R-Ras as a central regulator of reciprocal rewiring between ERK1/2 and ERK5 pathways under targeted MAPK inhibition. Functional disruption of this signaling circuit enhances melanoma cell death, providing a mechanistic rationale for co-targeting R-Ras together with MAPK signaling to limit adaptive responses to targeted therapy in BRAFV600E melanoma.

## INTRODUCTION

ERK5 and ERK1/2 are members of the Mitogen-Activated Protein Kinase (MAPK) family and play key roles in multiple physiological and pathological processes, including cancer (1). In melanoma, one of the most aggressive malignancies, recurrent mutations in BRAF (50-60%), N-Ras (20-25%), and NF1 (14%) drive constitutive activation of ERK1/2 pathway, promoting tumor onset and progression (2,3). The development of BRAF and MEK1/2 inhibitors has significantly improved the management of BRAFV600E-mutant melanoma. However, intrinsic or acquired resistance remains a major clinical challenge (4,5).

We recently showed that ERK5 inhibition reduces melanoma growth by promoting cell cycle arrest and cellular senescence (6,7). On the other hand, accumulating evidence in recent years points to the occurrence of ERK5 activation upon BRAF or MEK1/2 inhibition in both N- Ras- and BRAF-driven tumors in preclinical models. Consistently, combined targeting of ERK5 and the ERK1/2 pathway exerts greater antitumor activity than single-agent treatments (5,6,8–11). On the other hand, whether ERK5 inhibition triggers compensatory activation of the canonical RAF-MEK1/2-ERK1/2 pathway remains unknown.

We previously identified Ras-Related Protein (R-Ras), a small GTPase of the Ras family (12), among the most significantly upregulated genes following ERK5 depletion (7). R-Ras has been reported to interact with RAF proteins, Ral-GDS and is also able to activate the PI3K pathway (13–15). These observations prompted us to investigate whether R-Ras mediates the functional interplay between ERK5 and ERK1/2 signaling.

## MATERIALS AND METHODS

### Cells and cell culture

A375 melanoma cells (CVCL_0132) (16) were obtained from ATCC; SK-Mel-5 melanoma cells (CVCL_0527) (17) were kindly provided by Dr. Laura Poliseno (CRL-ISPRO, Pisa, Italy). HEK-239T cells (CVCL_0063) were obtained from ATCC. Cells were maintained in DMEM supplemented with 10% heat-inactivated fetal bovine serum (FBS), 2mmol/L glutamine, 50 U/mL penicillin and 50 mg/mL streptomycin (Euroclone). Cell lines were yearly authenticated by cell line profiling (Promega PowerPlex Fusion System Kit; BMR Genomics s.r.l.). The presence of mycoplasma was periodically tested by PCR.

### Drugs

ERK5 inhibitors (ERK5i) XMD8-92 (18) and JWG-071 (19) were from MedChemExpress LLC (Monmouth Junction, NJ, USA); BRAFi, vemurafenib (20), MEK1/2i, trametinib (21), ERK1/2i SCH772984 (22) were from Selleckchem (Selleckchem, Italy, www.selleckchem.com); pan-Rasi RMC-6236 (Daraxonrasib) (23) was obtained from Cayman Chemical (Ann Arbor, Michigan, USA).

### Cell lysis and Western Blot

Total cell lysates were obtained using Laemmli Buffer (24) or RIPA buffer (25) as reported previously. Proteins were separated by SDS-PAGE and transferred onto Amersham Protran® nitrocellulose membranes (GE healthcare) by electroblotting. Infrared imaging (Odissey, Li-Cor Bioscience) was performed. Images were quantified with ImageJ software. Antibodies used are listed in Table S1.

### Low-density phosphoprotein array

Low-density phosphoprotein arrays were performed with the PathScan® RTK Signaling Antibody Array Kit (#7949, Cell Signaling Technology, USA) following the manufacturer’s instructions. Total cell lysates were obtained using Cell Lysis Buffer (#9803, Cell Signaling Technology). Infrared imaging (Odissey, Li-Cor Bioscience) was performed. Images were quantified with ImageJ software.

### RNA interference and lentiviral vectors

Transfection of human melanoma cell lines was performed using the Lipofectamine™ 2000 (Invitrogen, Waltham, MA, USA) using smart Pool siRNA specific for human ERK5 or human R-Ras or non-targeting siRNA (sequence accession no. #NM_002749), as previously described (24) (Table S2). For single-gene silencing experiments, siRNAs were used at a final concentration of 100 nM. In double-silencing experiments, each siRNA was used at 50 nM, resulting in a total final siRNA concentration of 100 nM. siRNA specific for human ERK5 and non-targeting siRNA used were purchased from Dharmacon (Lafayette, USA) and siRNA for human R-Ras used were purchased from Sigma-Merck (St. Louis, MO, USA). Stable knockdown of ERK5 in melanoma cells was performed using lentiviral vectors. TRC1.5-pLKO.1-puro lentiviral vectors (Table S3) were produced in HEK-293T cells as reported previously (26). Transduced cells were selected with 2 µg/mL puromycin for at least 72 hours.

### Plasmids and Transfection

pCELF-RRASQL constructs were a kind gift from Mario Chiariello (CNR, Siena, Italy). pcDNA3.1-HA-ERK5wt construct was a kind gift from Atanasio Pandiella (CIC, Salamanca, Spain). HEK-293T cells were plated on six-well dishes (3 × 10^5^ cells/well) and transfected after 24 hours with a total amount of 3 μg of plasmid DNA using jetPEI® (Polyplus-transfection S.A, Illkirch, France), following the manufacturer’s instructions.

### Quantitative real-time PCR

Total RNA was isolated using TriFast II (Euroclone). cDNA synthesis was carried out using ImProm-II Reverse Transcription System (Promega Corporation, Madison, WI, USA), while quantitative PCR (qPCR) was performed using GoTaq qPCR Master Mix (Promega Corporation, Madison, WI, USA). Primer sequences are reported in Table S4.

PCR products were detected in the CFX96 Touch Real-Time PCR Detection System (Bio-Rad, Hercules, CA, USA). Results were analysed using CFX Maestro Software. A melting curve analysis was performed to discriminate between specific and non-specific PCR products. The relative expression of KLF2, MEF2C and R-Ras mRNA was calculated using a comparative threshold cycle method and the formula 2^-(DDCt)^ (27). The level of expression of mRNA of interest was normalized to that of both GAPDH and 18S mRNA.

### Flow cytometry analysis

Cell death (Annexin-V-FLUOS Staining Kit, no. 11988549001, Roche Diagnostics, Basel, Switzerland) was determined as previously reported using a FACSCanto (Beckton Dickinson) (28).

### Patient dataset

Expression of R-Ras in normal, primary, and metastatic tumor tissues was obtained from The Cancer Genome Atlas data set on TNMplot database (http://www.tnmplot.com) (29). Analysis of the relationship between R-Ras and p21 mRNA and R-Ras mRNA and RB1 PS807/S811 protein in melanoma patients was performed using the publicly available Skin Cutaneous Melanoma data set from The Cancer Genome Atlas (PanCancer Atlas) on cBioPortal for Cancer Genomics (https://www.cbioportal.org) (30,31).

### Statistical analysis

Statistical analysis was conducted via unpaired Student’s t test for comparisons between two groups or one-way analysis of variance (ANOVA), followed by the Bonferroni post hoc correction for multiple comparisons. P values less than 0.05 were considered statistically significant.

## RESULTS

### Activation of the MEK1/2-ERK1/2 axis upon ERK5 inhibition in BRAFV600E melanoma cells

We have recently shown that ERK5 inhibition reduces melanoma growth by blocking the cell cycle and inducing cellular senescence (6,7). To identify pro-proliferative pathways activated following ERK5 inhibition and that could reduce the impact of its targeting, we performed low-density phosphoprotein arrays. We found that both ERK5 knock-down (KD) using specific shRNA and pharmacological inhibition of either MEK5 (BIX02189) or ERK5 (XMD8-92) determine a marked increase in ERK1/2 phosphorylation in A375 melanoma cells (Fig. 1A). To substantiate these data, we performed Western Blot experiments using two metastatic melanoma cell lines harboring BRAFV600E mutation. Using this approach, we confirmed the increase of ERK1/2 phosphorylation in both A375 and Sk-Mel-5 cells after ERK5 KD with two different shRNAs (Fig. 1B). The same effects were observed following pharmacological ERK5 inhibition with XMD8-92 and the more specific ERK5i JWG-071 (Fig. 1C) (19). The increased ERK1/2 phosphorylation was accompanied by an increase of the activation of the ERK1/2 upstream kinase MEK1/2 (Fig. 1B, C). Effectiveness of ERK5 inhibition following pharmacological targeting was confirmed by the reduced mRNA levels of the downstream targets KLF2 and MEF2C (Fig. 1D, E) (32,33).

**Fig. 1.**
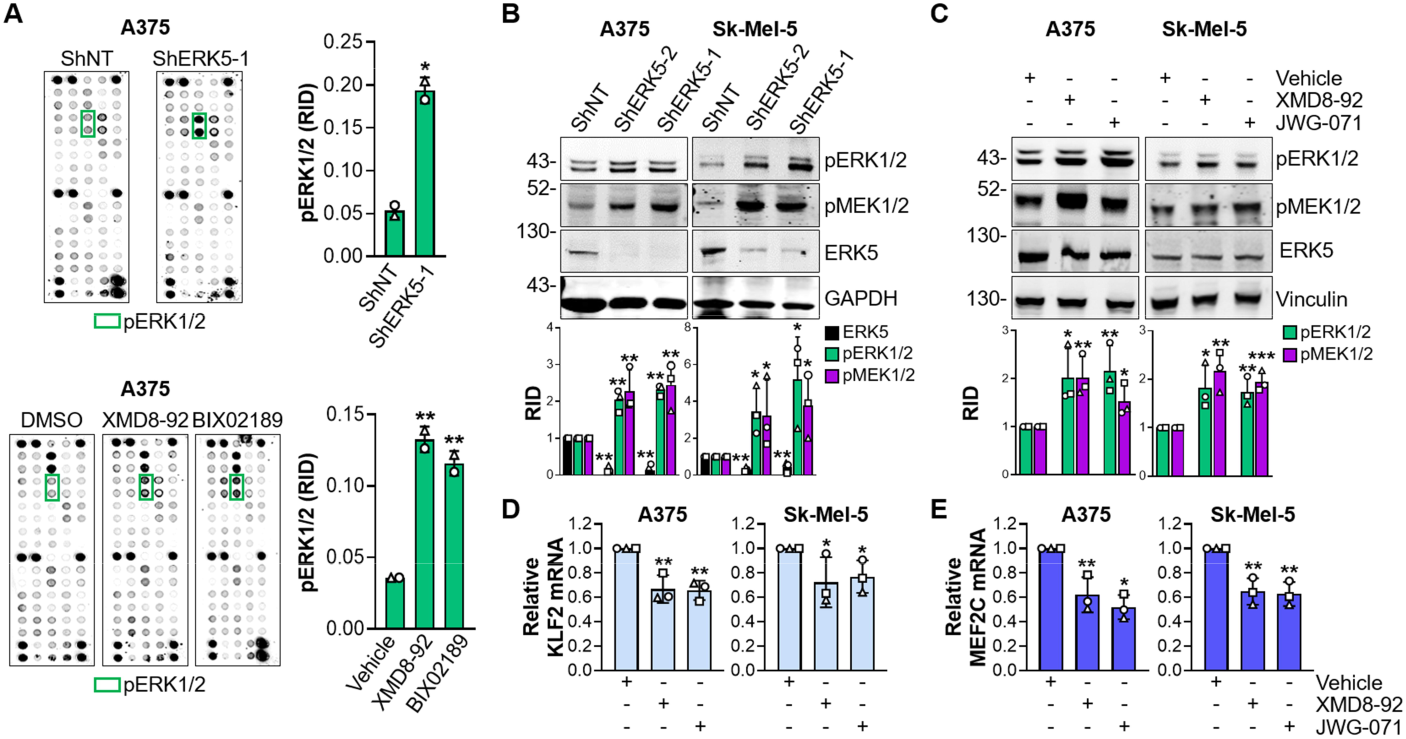
ERK5 inhibition elicits the increase of MEK1/2 and ERK1/2 phosphorylation. **A** A375 cells were transduced with lentiviral vectors carrying control non-targeting shRNA (shNT) or ERK5-specific shRNA (shERK5-1) and lysed after 5 days (top). A375 cells were treated with XMD8-92 (10 μM) or BIX02189 (10 μM) and lysed after 72 hours (bottom). Low-density phosphoprotein arrays were performed. Graphs show densitometric quantification of ERK1/2 phosphorylation as Relative Integrated Density (RID). Data, shown as mean ± SD, are from one experiment with two technical duplicates. **B** A375 and Sk-Mel-5 cells were infected with lentiviral vectors carrying shNT or two different shERK5 (1 and 2) and lysed after 5 days. Western Blot was then performed with the indicated antibodies. Migration of molecular weight marker is indicated on the left (kDa). Representative images of three independent experiments are shown. Graphs show the mean ± SD of the densitometric quantification of indicated proteins as RID. **C** A375 and Sk-Mel-5 cells were treated with DMSO (Vehicle), 5 μM XMD8-92 or 5 μM JWG-071 for 72 hours and lysed. Western Blot was then performed with the indicated antibodies. Migration of molecular weight marker is indicated on the left (kDa). Representative images from three independent experiments are shown. Graphs show the mean ± SD of the densitometric quantification of indicated proteins as RID. **D, E** Q-PCR analysis of KLF2 (D) and MEF2C (E) mRNA levels in A375 and Sk-Mel-5 cells following DMSO (Vehicle), 5 μM XMD8-92 or 5 μM JWG-071 for 72 hours. Graphs show mRNA levels from three independent experiments as mean ± SD. Statistical analyses were performed using two-tailed Student’s *t* test. \**P* < 0.05, ** *P* < 0.01 versus control (ShNT or Vehicle).

### ERK5 inhibition determines an increase of R-Ras mRNA and protein levels in BRAFV600E melanoma cells

To identify possible upstream regulators responsible for the increased MEK1/2 and ERK1/2 phosphorylation following ERK5 inhibition, we used data previously obtained from transcriptomic experiments performed in A375 and Sk-Mel-5 cells after ERK5 KD (7). Among the possible upstream activators, we found that R-Ras mRNA increased in both cell lines upon ERK5 KD (7). Here, we confirmed the increase of R-Ras mRNA (Fig. 2A) and protein levels (Fig. 2B) in lysates obtained from A375 and Sk-Mel-5 ERK5 KD cells with respect to controls. We also found a marked increase of R-Ras protein levels in both cell lines treated with two different ERK5 inhibitors (Fig. 2C). The efficacy of ERK5 inhibition was confirmed by increased p21 protein levels (7).

**Fig. 2.**
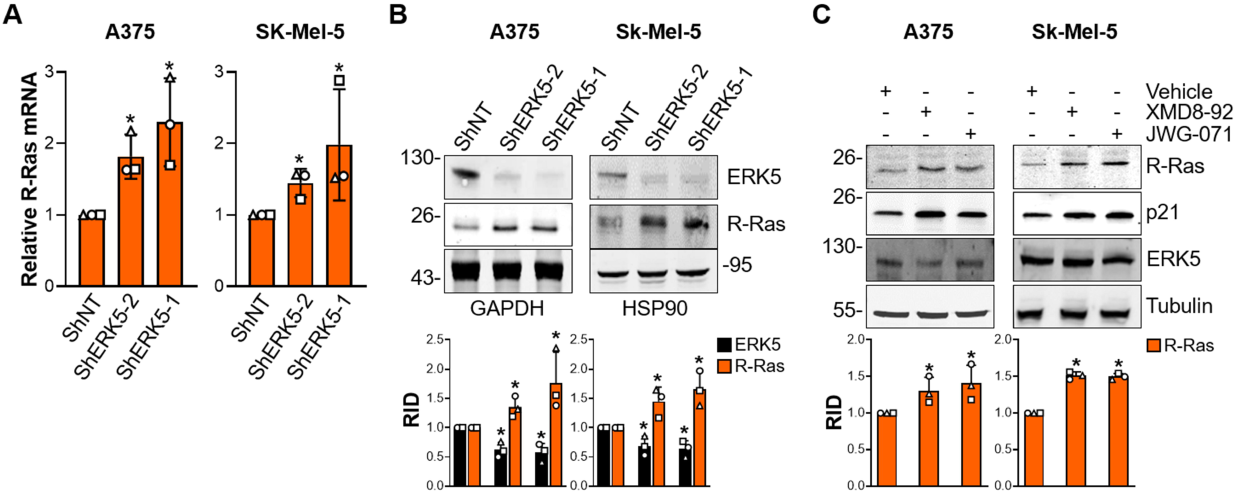
ERK5 inhibition elicits the increase of R-Ras mRNA and protein levels. **A** A375 and SK-Mel-5 cells were transduced with lentiviral vectors carrying control non-targeting shRNA (shNT) or ERK5-specific shRNA (shERK5-1 or -2). Five days after infection, cells were lysed and R-Ras mRNA levels were determined by Q-PCR. Graphs show mRNA levels from three independent experiments as mean ± SD. **B** A375 and Sk-Mel-5 cells were infected with lentiviral vectors carrying shNT or shERK5 (shERK5-1 or -2) and lysed after 5 days. Western Blot was then performed with the indicated antibodies. Migration of molecular weight marker is indicated on the left (kDa). Representative images of three independent experiments.are shown. Graphs show the mean ± SD of the densitometric quantification of indicated proteins as RID. **C** A375 and Sk-Mel-5 cells were treated with DMSO (Vehicle), 5 μM XMD8-92 or 5 μM JWG-071 for 72 hours and lysed. Western Blot was then performed with the indicated antibodies. Migration of molecular weight marker is indicated on the left (kDa). Representative images of three independent experiments are shown. Graphs show the mean ± SD of the densitometric quantification of indicated proteins as RID. Statistical analyses were performed using two-tailed Student’s *t* test. \**P* < 0.05 versus control (ShNT or Vehicle).

### R-Ras mRNA levels in melanoma patients

The publicly available Skin Cutaneous Melanoma dataset from the Cancer Genome Atlas (TCGA) on TNMplot provided evidence that the mRNA levels of R-Ras are lower in primary and metastatic melanomas than in healty control tissues (Fig. 3A). These results are in line with a recent report showing that higher R-Ras mRNA levels are associated with improved overall survival in the same TCGA melanoma cohort (34). Furthermore, analysis of the same TCGA cohort through cBioPortal revealed a positive correlation between R-Ras and p21 mRNA levels (Fig. 3B), together with a negative correlation between R-Ras mRNA expression and RB1 phosphorylation at Ser807/Ser811 (Fig. 3C). Collectively, these observations support a potential tumor-suppressive role for R-Ras in melanoma.

**Fig. 3.**
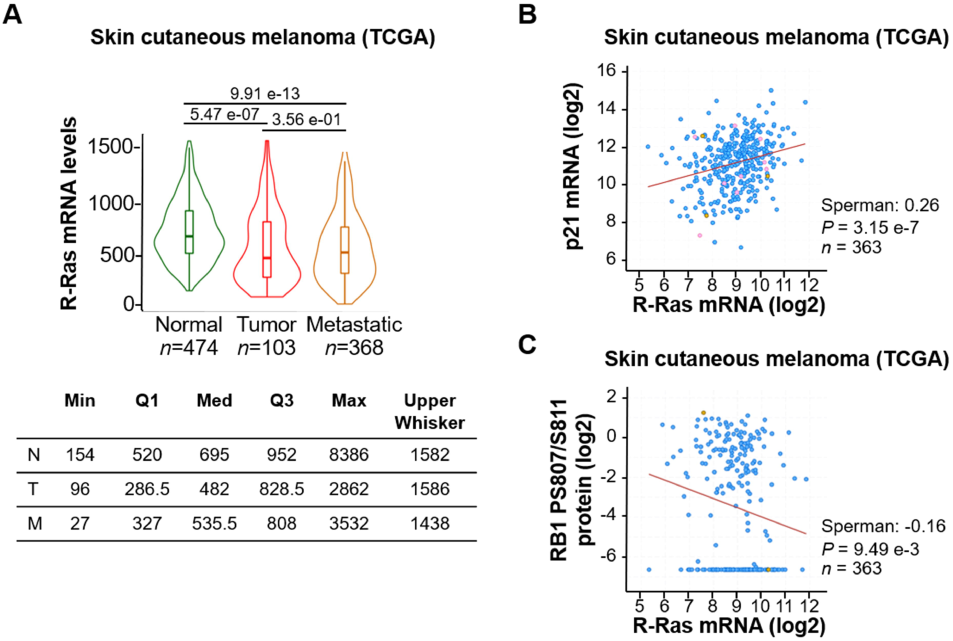
Clinical impact of R-Ras expression levels in melanoma patients. **A** Violin plots show R-Ras gene expression profile in normal skin (Normal), and primary (Tumor) or metastatic (Metastatic) melanoma obtained by the Skin Cutaneous Melanoma (SKCM) data set (The Cancer Genome Atlas, TCGA) on TNMplot. *P* values were computed using Dunnett’s test. **B**, **C** Expression levels of R-Ras and p21 mRNA (B) and expression levels of R-Ras mRNA and RB1 PS807/S811 protein (C) from the Skin Cutaneous Melanoma (SKCM) data set [The Cancer Genome Atlas (TCGA)] on cBioPortal.

### R-Ras knock down reverts ERK1/2 activation and potentiates the cytotoxicity upon ERK5 inhibition in BRAFV600E melanoma cell lines

To deepen the possible involvement of R-Ras in ERK1/2 activation following the inhibition of ERK5, R-Ras KD was performed alone or in combination with pharmacological inhibition of ERK5 using JWG-071. In both cell lines, R-Ras KD using specific siRNA prevented the increase of ERK1/2 phosphorylation observed upon JWG-071 treatment (Fig. 4A). These results support an involvement of R-Ras in ERK1/2 activation induced by the inhibition of the ERK5 pathway.

**Fig. 4.**
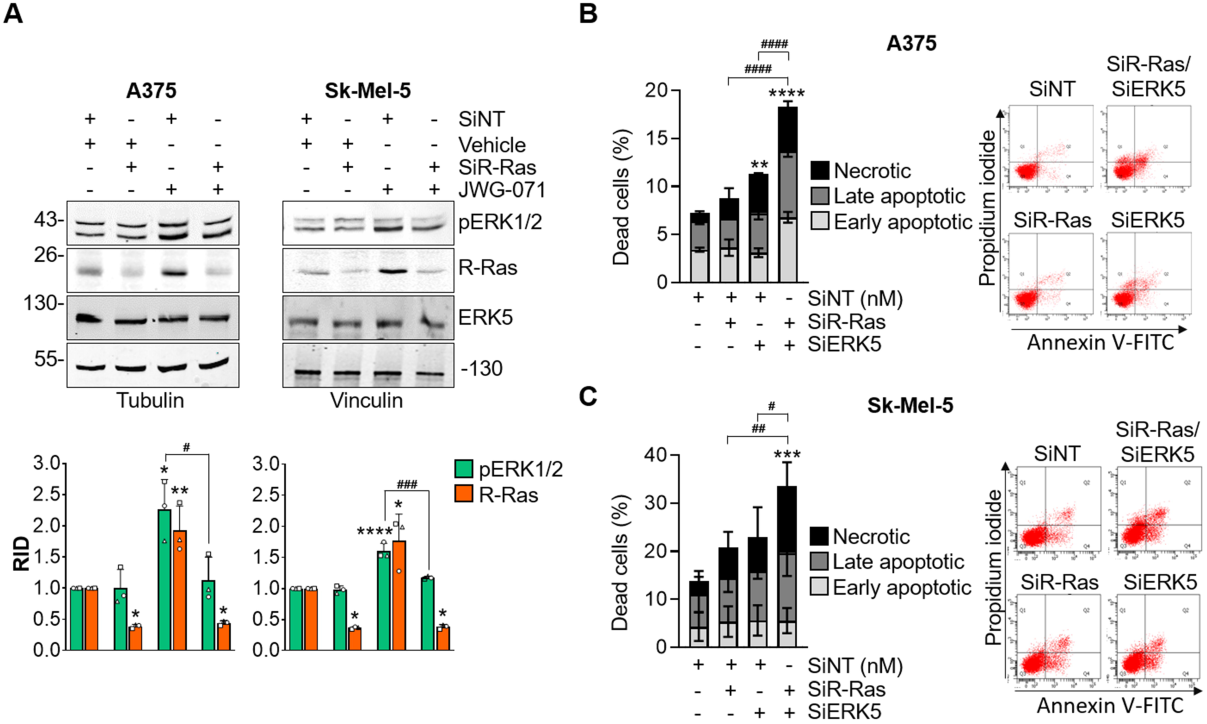
R-Ras KD prevents ERK1/2 activation elicited by ERK5 inhibition and potentiates the cytotoxic effect of ERK5 inhibition. **A** A375 and Sk-Mel-5 cells were transfected with non-targeting siRNA (siNT) or R-Ras specific siRNA (siR-Ras) and treated with DMSO (Vehicle) or 5 µM JWG-071 for 72 hours before lysis. Western Blot was then performed with the indicated antibodies. Migration of molecular weight marker is indicated on the left (kDa). Representative images from three independent experiments are shown. Graphs show the mean ± SD of the densitometric quantification of indicated proteins as RID. **B**, **C** Annexin V/Propidium Iodide assay was performed to quantify dead cells. A375 (B) and Sk-Mel-5 (C) cells were transfected with non-targeting siRNA (siNT), R-Ras (siR-Ras) or ERK5 (siERK5) targeting siRNA for 72 hours before lysis. The graphs show the percentage of dead cells. Data, shown as mean ± SD, are from three independent experiments. Representative dot plot for each condition is shown on the right. Statistical analyses were performed using one-way ANOVA test. \**P* < 0.05, \*\**P* < 0.01, \*\*\**P* < 0.001, \*\*\*\**P* < 0.0001 versus control (SiNT+Vehicle or SiNT). ^#^*P* < 0.05, ^##^*P* < 0.01, ^###^*P* < 0.001, ^####^*P* < 0.0001 between indicated conditions.

To understand whether R-Ras-mediated ERK1/2 activation following ERK5 inhibition exerts a positive or negative role in supporting cancer cell survival, cell death was assessed by Annexin V/propidium iodide staining. (Fig. 4B, C). Combined R-Ras and ERK5 KD, using specific siRNA, significantly increased the percentage of dead cells in both A375 and Sk-Mel-5 cells compared with either single treatment (Fig. 4B, C). Similar effects were observed when R-Ras KD was combined with pharmacological ERK5 inhibition using JWG-071 (Fig. 5A, B). Consistently, combined R-Ras KD and JWG-071 treatment resulted in a significantly greater reduction in melanoma cell number compared with either condition alone (Fig. 5C, D).

**Fig. 5.**
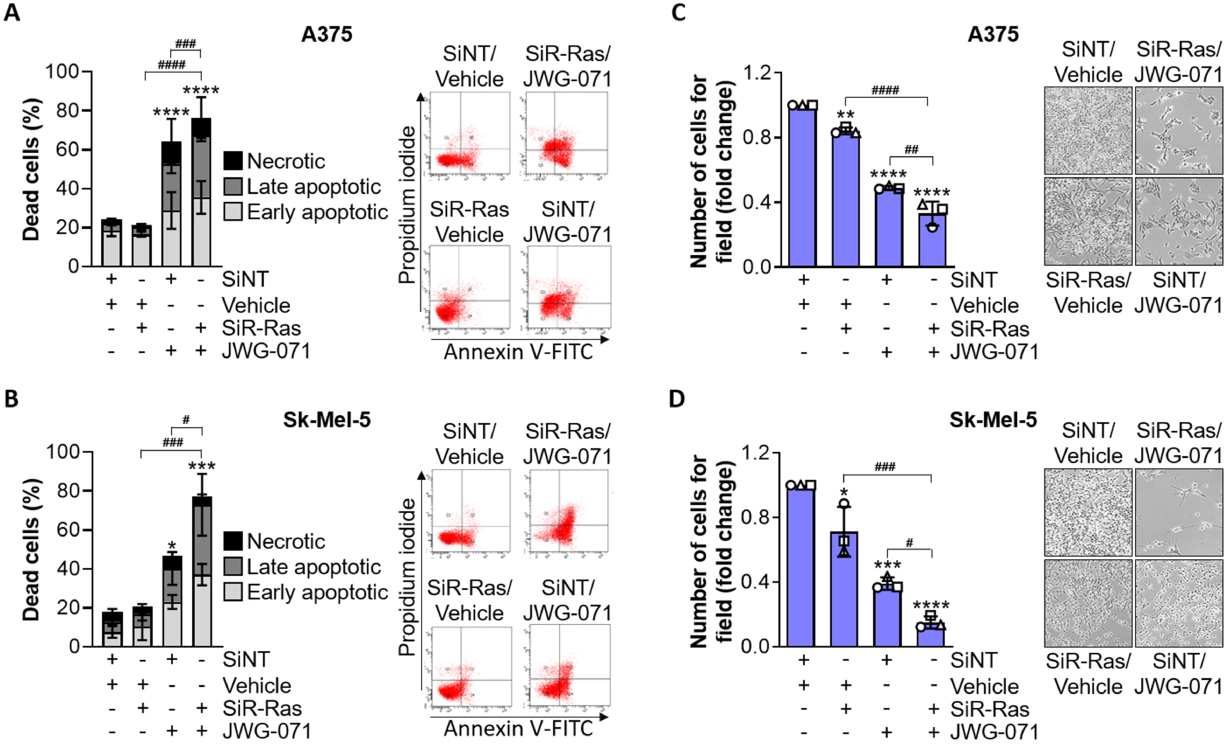
R-Ras KD potentiates the cytotoxic effect of ERK5 inhibition. **A, B** A375 (**A**) or Sk-Mel-5 (**B**) cells were transfected with non-targeting siRNA (siNT) or R-Ras specific siRNA (siR-Ras) and treated with DMSO (Vehicle) or 5 µM JWG-071 for 72 hours before lysis. Annexin V/Propidium Iodide assay was performed to quantify dead cells. Graphs show the percentage of dead cells as mean ± SD from three independent experiments. Representative dot plot for each condition is shown. **C**, **D** A375 (C) or SK-Mel-5 (D) cells were treated as above. Viable cell quantification was performed by counting cells in six random images/well, taken using brightfield microscope 20X. One representative image for each experimental condition is shown. Data, shown as mean ± SD, are from three independent experiments. Statistical analyses were performed using one-way ANOVA test. \**P* < 0.05, \*\**P* < 0.01, \*\*\**P* < 0.001, \*\*\*\**P* < 0.0001 versus control (siNT + Vehicle). ^#^*P* < 0.05, ^##^*P* < 0.01, ^###^*P* < 0.001, ^####^*P* < 0.0001 between indicated conditions.

Collectively, these findings indicate that R-Ras promotes melanoma cell survival following ERK5 inhibition, likely by mediating compensatory pro-survival signaling pathways, including the activation of the MEK1/2-ERK1/2 cascade.

### R-Ras is involved in ERK5 activation upon Raf-MEK1/2-ERK1/2 targeting in BRAFV600E melanoma cells

Accumulating evidence indicates that inhibition of the RAF-MEK1/2-ERK1/2 pathway leads to the activation of ERK5 (5) in preclinical cancer models. We confirmed this effect in both A375 and Sk-Mel-5 cells after treatment with SCH779284 (ERK1/2i) or vemurafenib (BRAFV600Ei) in combination with trametinib (MEK1/2i) (Fig. 6A). A significant increase of ERK5 phosphorylation at MEK5-consensus sites was observed in both treatment conditions compared to the controls. Intriguingly, increased ERK5 activation was accompanied by an increase in R-Ras protein levels. Effectiveness of RAF-MEK1/2-ERK1/2 inhibiting treatments was confirmed by the reduced phosphorylation of ERK1/2. Increased ERK5 activation was also confirmed by the higher KLF2 mRNA levels (Fig. 6B). Notably, R-Ras mRNA levels were increased following RAF-MEK1/2-ERK1/2 targeting (Fig. 6C).

**Fig. 6.**
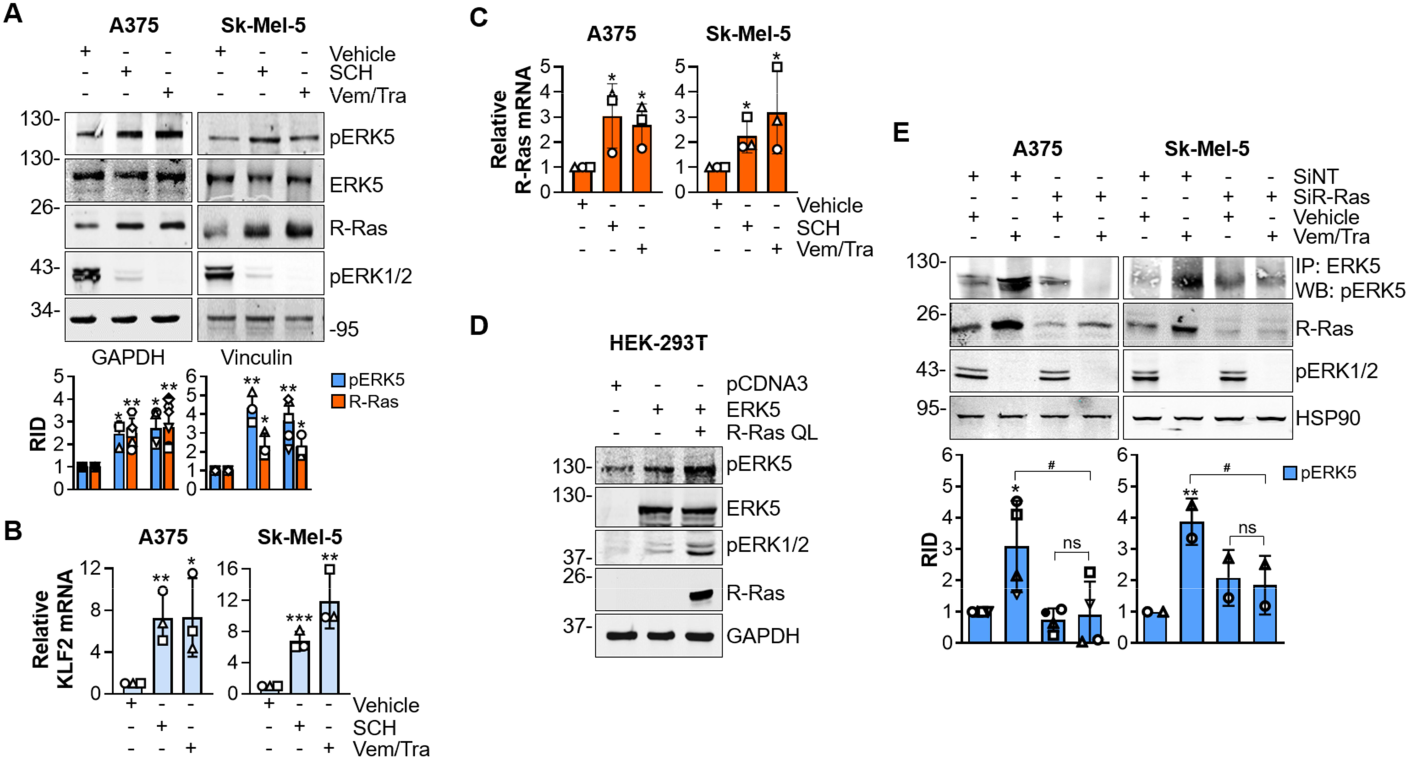
R-Ras KD prevents ERK5 phosphorylation induced by the BRAF-MEK1/2-ERK1/2 pathway inhibition. **A** A375 and Sk-Mel-5 cells were treated with DMSO (Vehicle), 1 μM SCH772984 or 1 μM vemurafenib/1μM trametinib for 72 hours and lysed. Western Blot was then performed with the indicated antibodies. Migration of molecular weight marker is indicated on the left (kDa). Representative images from three independent experiments are shown. Graphs show the mean ± SD of the densitometric quantification of indicated proteins as RID. **B, C** Q-PCR analysis of KLF2 (B) and R-Ras (C) mRNA levels in A375 and Sk-Mel-5 cells following DMSO (Vehicle), 1 μM SCH772984 or 1 μM vemurafenib/1μM trametinib for 72 hours. Graphs show mRNA levels from three independent experiments as mean ± SD. **D** HEK-293T cells were transfected with equimolar amounts of pcDNA3 (control) or wt ERK5 in combination with pcDNA3 or mutant active R-Ras (R-Ras QL) plasmids for 24 hours before lysis. Western Blot was then performed with the indicated antibodies. Migration of molecular weight marker is indicated on the left (kDa). Representative images from two independent experiments. **E** A375 and Sk-Mel-5 cells were transfected with non-targeting siRNA (siNT) or R-Ras specific siRNA (siR-Ras) for 72 hours and then treated with DMSO (Vehicle) or 1μM vemurafenib/1μM trametinib for further 24 hours before lysis. The levels of ERK5 phosphorylation was evaluated following immunoprecipitation. Western Blot was then performed with the indicated antibodies. Migration of molecular weight marker is indicated on the left (kDa). Representative images from three (A375) or two (Sk-Mel-5) independent experiments are shown. Graphs show the mean ± SD of the densitometric quantification of indicated proteins as RID. Statistical analyses were performed using appropriate tests based on data distribution: two-sided unpaired Student’s t test was applied for (**A, B, C, E** Sk-Mel-5), and one-way ANOVA test was used for (**E** A375). \**P* < 0.05, ** *P* < 0.01, \*\*\**P* < 0.001 versus control (Vehicle or siNT + Vehicle). ^#^*P* < 0.05 between indicated conditions. ns, not significant.

The contribution of R-Ras to ERK1/2 signaling remains controversial (35–37) while its role in the ERK5 pathway has never been addressed. To test wether R-Ras is among the upstrem activators of both ERK1/2 and ERK5, we overexpressed a constitutive active form of R-Ras (R-Ras QL) in HEK-293T cells (Fig. 6D). Overexpression of constitutively active R-Ras enhanced the phosphorylation of either ERK1/2 or of ectopic ERK5, positioning R-Ras upstream both MAPK. The possible involvement of R-Ras in ERK5 activation following ERK1/2 pathway inhibition was then investigated (Fig. 6E). In both A375 and Sk-Mel-5 cells, R-Ras KD impaired the increase of ERK5 phosphorylation induced by vemurafenib and trametinib combination, pointing to an involvement of R-Ras in ERK5 activation in these experimental conditions.

To understand whether R-Ras-dependent ERK5 activation is a resistance strategy in response to suppression of ERK1/2 pathway, combined inhibition of the ERK1/2 pathway and of R-Ras was performed to evaluate cell death in A375 and Sk-Mel-5 cells (Fig. 7). Combined genetic inhibition of R-Ras and pharmacological inhibition of Raf-MEK1/2-ERK1/2 pathway exceeded the effects of each treatment alone, in both A375 (Fig. 7A) and Sk-Mel-5 (Fig. 7B) cell lines.

**Fig. 7.**
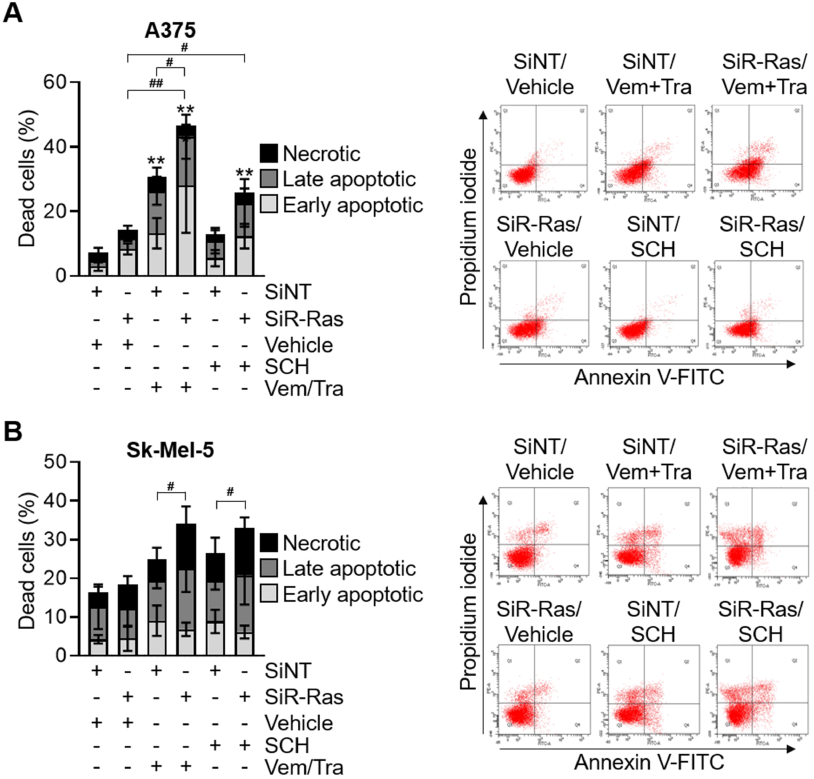
R-Ras KD potentiates the cytotoxic effect of BRAF-MEK1/2-ERK1/2 inhibition. **A**, **B** A375 (A) and Sk-Mel-5 (B) cells were transfected with non-targeting siRNA (siNT) or R-Ras-specific siRNA (siR-Ras) for 72 hours and then treated with DMSO (Vehicle) or 1 μM vemurafenib/1 μM trametinib or 1 μM SCH772984 for a further 24 hours. Annexin V/Propidium Iodide assay was performed to quantify dead cells. The graphs show the percentage of dead cells. Data, shown as mean ± SD, are from three independent experiments. Representative dot plot for each condition is shown. Statistical analyses were performed using one-way ANOVA test was used. \*\**P* < 0.01 versus control (siNT + Vehicle). ^#^*P* < 0.05, ^##^*P* < 0.01 between indicated conditions.

### The pan-Ras inhibitor RMC-6236 potentiates the antiproliferative effects of ERK5 or MAPK-ERK pathway inhibition in BRAFV600E melanoma cells

To evaluate the translational potential of these findings, we tested the effects of the pan-Ras inhibitor RMC-6236 on the observed reciprocal MAPK rewiring following targeted inhibition pressure (23,35,38). As few lines of evidence have included R-Ras among the targets of RMC-6236 (35), we tested the efficacy of this drug by overexpressing R-RAS QL in HEK-293T cells (Fig. 8A), and found that treatment with RMC-6236 prevented R-RAS QL-induced ERK1/2 activation. RMC-6236 treatment decreased the number of viable cells in a dose-dependent manner in both A375 and Sk-Mel-5 cell lines (Fig. 8B). Of note, IC values were relatively higher than those observed in N-Ras mutated melanoma cell lines (39). The combination of RMC-6236, used at around IC50 concentrations, with vemurafenib+tramentinib or SCH779284 or JWG-071 robustly impacted cell viability of A375 and Sk-Mel-5 cells (Fig. 8C), with respect to single treatments.

**Fig. 8.**
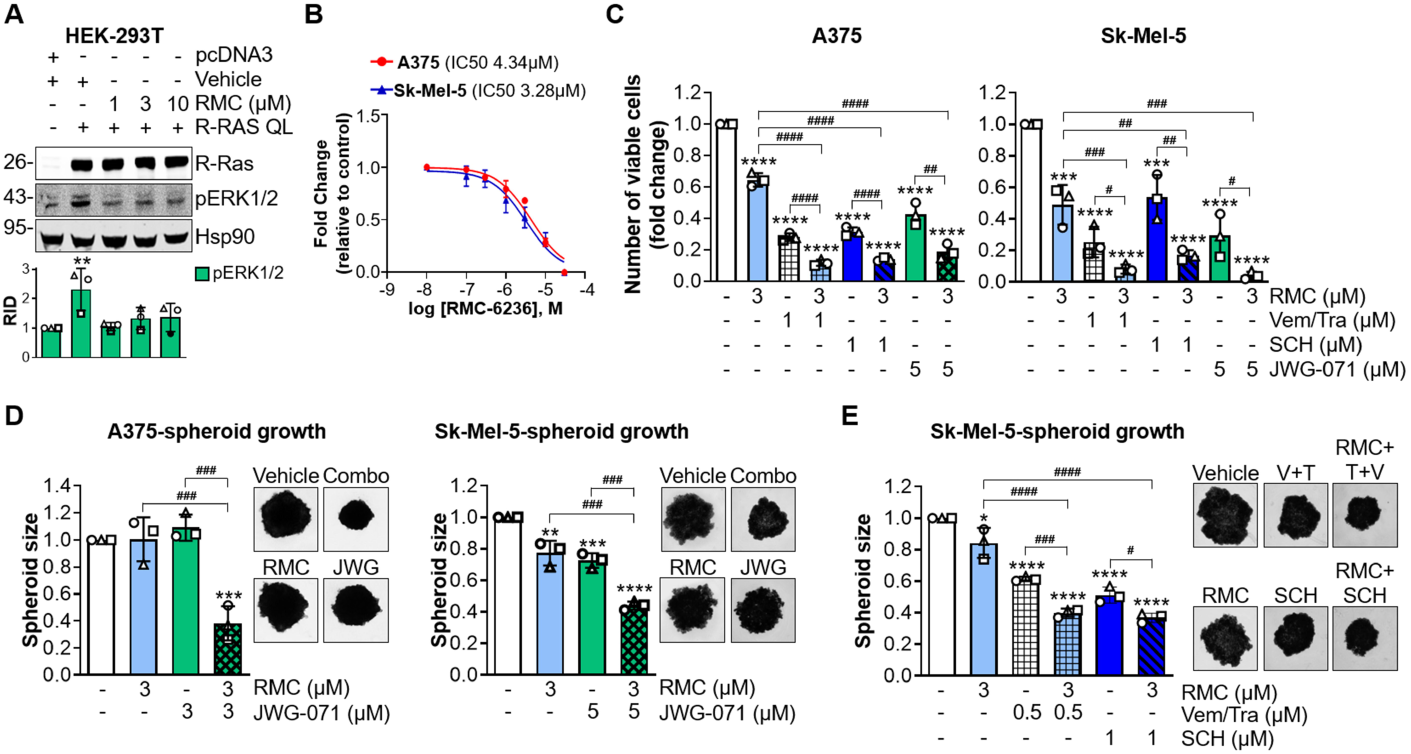
Ras pharmacological inhibition potentiates the antiproliferative effects of either ERK5 or ERK1/2 pathway inhibition. **A** HEK-293T cells were transfected with equimolar amounts of pcDNA3 (control) or mutant active R-Ras (R-Ras QL) plasmids for 24 hours and treated for a further 4 hours with DMSO (Vehicle) or RMC-6236 at the indicated concentration before lysis. Western Blot was then performed with the indicated antibodies. Migration of molecular weight marker is indicated on the left (kDa). Representative images are shown. Graphs show densitometric quantification of ERK1/2 phosphorylation as Relative Integrated Density (RID). Data, shown as mean ± SD, are from three independent experiments. **B** Dose-response curves of RMC-6236 in A375 or Sk-Mel-5 cells after 72 hours of treatment or DMSO (Vehicle). Viability was assessed by Trypan blue exclusion assay to determine IC50 values. Data, shown as mean ± SD, are from three independent experiments. **C** A375 and Sk-Mel-5 cells were treated with DMSO (-), 3 μM RMC-6236 (RMC), 1 μM vemurafenib/1 μM trametinib (Vem/Tra), 1 μM SCH722984 (SCH) or 5 μM JWG-071 alone or in combination for 72 hours. Viable cell number was then determined by Trypan blue exclusion. Data, shown as mean ± SD, are from three independent experiments. **D** A375 and Sk-Mel-5 spheroids were treated with DMSO (-), 3 μM RMC-6236 (RMC), 3 (A375) or 5 (Sk-Mel-5) μM JWG-071 alone or in combination. Graphs shows the quantification of spheroid volume after 5 days normalized for the time point 0. Data, shown as mean ± SD, are from three independent experiments. Representative images of spheroids taken at day 5 are shown. **E** Sk-Mel-5 spheroids were treated with DMSO (-), 3 μM RMC-6236 (RMC), 0.5 μM vemurafenib/0.5 μM trametinib (Vem/Tra) or 1 μM SCH722984 (SCH) alone or in combination. Graphs shows the quantification of spheroid volume after 5 days normalized for the time point 0. Data, shown as mean ± SD, are from three independent experiments. Representative images of spheroids taken at day 5 are shown. Statistical analyses were performed using one-way ANOVA test. \**P* < 0.05, \*\**P* < 0.01, \*\*\**P* < 0.001, \*\*\*\**P* < 0.0001 versus control (pcDNA3 + Vehicle or Vehicle). ^#^*P* < 0.05, ^##^*P* < 0.01, ^###^*P* < 0.001, ^####^*P* < 0.0001 between indicated conditions.

We then used A375 and Sk-Mel-5 spheroids to move to in vitro 3D models of tumour growth. The combination of RMC-6236 and JWG-071, at concentrations determining no (A375) or slight (Sk-Mel-5) effects when used alone in 3D cultures, resulted in a drop of more than 50% of melanoma spheroid volumes (Fig. 8D). Likewise, combined treatment with RMC-6236 and BRAF-MEK1/2-ERK1/2 inhibitors was more effective than single treatments in reducing 3D growth of Sk-Mel-5 cells (Fig. 8E). Collectively, these results indicate that the effects of RMC-6236 and ERK5 or BRAF-MEK1/2-ERK1/2 inhibitors on reducing cancer cell proliferation closely mirror those observed upon combined R-Ras KD and ERK5 or BRAF-MEK1/2-ERK1/2 inhibitors, supporting the potential translation of this strategy to the clinic.

## DISCUSSION

Advanced melanoma remains one of the most aggressive malignancies due to its remarkable molecular plasticity, which underlies intrinsic and acquired resistance to current therapies. Elucidating the mechanisms that sustain melanoma cell survival and adaptive responses is therefore critical to identify novel strategies to overcome therapeutic resistance. The clinical implementation of BRAF and MEK1/2 inhibitors has significantly improved the management of BRAFV600E-mutant melanoma. However, the long-term efficacy of these targeted therapies is frequently limited by compensatory signaling mechanisms that drive adaptive resistance. Accumulating evidence obtained in preclinical models points to ERK5 activation as a compensatory mechanism occurring upon RAF-MEK1/2-ERK1/2 inhibition, thereby promoting resistance to these therapeutic strategies (5). However, whether the reverse also occurs, namely if ERK5 inhibition elicits compensatory activation of ERK1/2 signaling, has remained unknown. Here, we identify a novel bidirectional crosstalk between the two MAPK pathways and demonstrate that the small GTPase R-Ras acts as a critical mediator of an adaptive response following targeted therapy in BRAFV600E-mutant melanoma cells. Our data show that genetic or pharmacological inhibition of ERK5 further increases MEK1/2 and ERK1/2 phosphorylation, which are constitutively high as a consequence of mutant BRAF, together with an increase in R-Ras mRNA and protein levels. Importantly, R-Ras silencing completely prevents this hyperactivation, demonstrating that ERK5 inhibition triggers a R-Ras-dependent compensatory hyperactivation of the canonical MAPK cascade. Although ERK5 inhibition alone impairs melanoma cell proliferation (6,7), the superior efficacy of combined ERK5 and R-Ras targeting in reducing cell viability indicates that R-Ras-driven ERK1/2 hyperactivation represents an adaptive feedback mechanism that promotes melanoma cell survival upon ERK5 inhibition.

Another intriguing finding of this work is that R-Ras is also involved in the previously reported activation of ERK5 following the inhibition of the BRAF-MEK1/2-ERK1/2 pathway (5). Indeed, either BRAFV600E+MEK1/2 (vemurafenib+trametinib) or ERK1/2 (SCH-772984) targeting induces ERK5 activation/phosphorylation together with R-Ras upregulation, whereas R-Ras silencing abolishes ERK5 phosphorylation and significantly enhances cell death.

Our data, therefore, identify R-Ras as a common upstream regulator of compensatory signaling in both directions, allowing melanoma cells to switch between ERK1/2 and ERK5 pathways following single-pathway inhibition. This reciprocal signaling network provides a mechanistic explanation for the enhanced efficacy observed upon combined pathway targeting and highlights MAPK network plasticity as a key determinant of therapeutic adaptation.

In silico analysis of melanoma patient datasets shows that R-Ras expression is lower in primary and metastatic tumors compared with normal skin, in line with a recent report showing that higher R-Ras levels correlate with improved overall survival (34). Moreover, we provide evidence that R-Ras expression positively correlates with p21 mRNA and negatively correlates with phosphorylated RB in melanoma patient datasets. These lines of evidence support the notion that higher R-Ras levels are linked to a less proliferative cellular state and suggest a potential role of R-Ras in restraining proliferative programs. However, whether this reflects a causal role of R-Ras in enforcing cell-cycle arrest or a broader transcriptional program associated with reduced tumor aggressiveness in which R-Ras is among the many players remains to be determined. Rather than being contradictory, these findings likely reflect distinct biological states. Basal R-Ras expression in untreated melanomas may be associated with a less proliferative tumor phenotype, whereas therapy-induced R-Ras upregulation represents an adaptive response that enables melanoma cells to maintain MAPK signaling and survive under pharmacological pressure.

Direct targeting of oncogenic Ras has recently become feasible, either through KRAS-selective inhibitors or emerging pan-Ras approaches (40,41). As for the latter, the pan-Ras inhibitor RMC-6236, used in this study, is currently being evaluated in a Phase III clinical trial for patients with previously treated RAS-mutated non-small cell lung cancer (NCT06881784). Moreover, RMC-7977, from which RMC-6236 have been obtained, showed strong antitumor activity in NRAS-mutant melanoma models, driving rapid regression via MAPK inhibition and immune activation (39). Here, we show that RMC-6236 effectively suppresses R-Ras-driven MAPK rewiring following therapeutic pressure, mirroring the effects of R-Ras knockdown when combined with either ERK5 or BRAF-MEK1/2-ERK1/2 inhibitors. In 3D melanoma models, these combinations markedly reduce tumor growth compared with single-agent treatments, supporting the potential clinical relevance of co-targeting R-Ras and MAPK pathways.

Mechanistically, our findings identify R-Ras as a nodal regulator of MAPK network plasticity in BRAFV600E-mutant melanoma. By enabling compensatory activation of ERK1/2 following ERK5 inhibition, and of ERK5 following ERK1/2 inhibition, R-Ras allows melanoma cells to preserve proliferative and survival signaling under targeted therapeutic pressure. Future studies should define the molecular mechanisms responsible for increased R-Ras expression under ERK5 or ERK1/2 blockade, including gene expression remodelling at R-Ras promoter and post-transcriptional regulation.

Finally, the development of selective and clinical grade R-Ras inhibitors, together with the identification of predictive biomarkers of response, will be essential for translating these findings into therapeutic strategies. In particular, determining whether R-Ras expression or activation status correlates with response to MAPK-targeted therapies may help guide patient stratification and rational combination treatments.

In conclusion, we uncovered a novel R-Ras-dependent reciprocal activation mechanism between ERK1/2 and ERK5 pathways in BRAFV600E-mutant melanoma, providing mechanistic insight into adaptive resistance to MAPK-targeted therapies. Therapeutic strategies that combine ERK5 or ERK1/2 inhibition with R-Ras targeting represent a promising approach to reduce tumor burden and to limit compensatory prosurvival signals, thus improving the efficacy of current melanoma treatments.

## ACKNOWLEDGEMENTS

The research leading to these results has received funding from AIRC (IG 2018 - ID. 21349 project and IG 2025 - ID. 32014 project - P.I. ER), Fondazione CR Firenze, European Union-NextGenerationEU-National Recovery and Resilience Plan, Mission 4 Component 2-Investment 1.5-THE-Tuscany Health Ecosystem-ECS00000017-CUP B83C22003920001 and from Fondo Beneficenza Intesa San Paolo (to ER). AM PhD fellowship was supported in part by *Fondazione BENEFICENTIA Stiftung* (to ER). YS PhD fellowship is supported by NRRP (D.M. 118/2023)/NEXTGENERATIONEU (to ER).

