## Supplementary Table 1 for "R-Ras coordinates reciprocal activation of ERK5 and ERK1/2 under single pathway inhibition in melanoma"

**Table S1.** List of the antibodies used in the study.

| ERK5 | Rabbit polyclonal | #3372 | Cell Signaling Technology, USA |
| --- | --- | --- | --- |
| ERK5 (C-7) | Mouse monoclonal | sc-398015 | Santa Cruz Biotechnology, USA |
| GAPDH | Mouse monoclonal | sc-47724 | Santa Cruz Biotechnology, USA |
| HSP90 | Mouse monoclonal | sc-13119 | Santa Cruz Biotechnology, USA |
| pERK1/2 (Thr202/Tyr204) | Rabbit polyclonal | #9101 | Cell Signaling Technology, USA |
| pERK5 (Thr218/Tyr220) | Rabbit polyclonal | #3371 | Cell Signaling Technology, USA |
| pMEK1/2 (Ser217/221) | Rabbit polyclonal | #9121 | Cell Signaling Technology, USA |
| p21 | Rabbit monoclonal | #2947 | Cell Signaling Technology, USA |
| R-Ras | Rabbit polyclonal | #8446 | Cell Signaling Technology, USA |
| Vinculin | Mouse monoclonal | V9131 | Merck, Germany |
| α-Tubulin | Mouse monoclonal | sc-32293 | Santa Cruz Biotechnology, USA |
