## Supplementary Table 2 for "R-Ras coordinates reciprocal activation of ERK5 and ERK1/2 under single pathway inhibition in melanoma"

**Supplementary Table S2**. List and sequences of the siRNAs used in the present study.

| **siRNA** | **Catalog #** | **Sense sequence** |
| --- | --- | --- |
| Non-targeting control | D-001210-01 | 5’-UAGCGACUAAACACAUCAAUU-3’ |
| ERK5 | M-003513-02 | 5’-CAUGAACCCUGCCGAUAUUUU-3’  5’-GCCCAGCGCUCGCAUCUCAUU-3’  5’-GAACUGUGAGCUCAAGAUUUU-3’  5’-AAACCAGUCUUUCGACAUGUU-3’ |
| R-RAS | SASI_HS01_00109592 | 5’-CGCAGAUUCUGCGGGUCAA-3’ |
