## Supplementary Table 3 for "R-Ras coordinates reciprocal activation of ERK5 and ERK1/2 under single pathway inhibition in melanoma"

**Supplementary Table S3**. List and sequences of the shRNAs used in the present study.

| **Gene** | **Clone** | **shRNA** | **TRC number** | **Sense sequence 5’ to 3’** |
| --- | --- | --- | --- | --- |
| none | none | shNT | TRCN0000023236 | CCGGCGACAATATCATCGCCATCAACTCGAGTTGATGGCGATGATATTGTCGTTTTT |
| MAPK7 | NM_139032 | shERK5-1 | TRCN0000010275 | CCGGGCCAAGTACCATGATCCTGATCTCGAGATCAGGATCATGGTACTTGGCTTTTT |
| MAPK7 | NM_139032 | shERK5-2 | TRCN0000010262 | CCGGGCTGCCCTGCTCAAGTCTTTGCTCGAGCAAAGACTTGAGCAGGGCAGCTTTTT |
