## Supplementary Table 4 for "R-Ras coordinates reciprocal activation of ERK5 and ERK1/2 under single pathway inhibition in melanoma"

**Supplementary Table S4.** List and sequences of the primers used for Q-PCR.

| **Gene** | **Primer sequence (5’ to 3’)** |
| --- | --- |
| KLF2 | Forward: CCAAGAGTTCGCATCTGAAGGC  Reverse: CCGTGTGCTTTCGGTAGTGGC |
| MEF2C | Forward: TCCACCAGGCAGCAAGAATACG  Reverse: GGAGTTGCTACGGAAACCACTG |
| RRAS | Forward: CTGCTGGTGTTCGCCATTAACG  Reverse: GATCTGCCTTGTTCCCGACCAA |
| GAPDH | Forward: AACAGCCTCAAGATCATCAGCAA  Reverse: CAGTCTGGGTGGCAGTGAT |
| 18s rRNA | Forward: CGGTACCACATCCAAGGAA  Reverse: GCTGGAATTACCGCGGCT |
